# High-titer production of aromatic amines in metabolically engineered *Escherichia coli*

**DOI:** 10.1101/2022.01.14.476304

**Authors:** Taiwei Yang, Peiling Wu, Yang Zhang, Jifeng Yuan

## Abstract

Aromatic amines are widely used in the pharmaceutical industry. Here, we reported the establishment of a bacterial platform for synthesizing three types of aromatic amines, namely, tyramine, dopamine, and phenylethylamine. Firstly, we expressed aromatic amino acid decarboxylase from *Enterococcus faecium* (*pheDC*) in an *Escherichia coli* strain with an increased shikimate (SHK) pathway flux toward L-tyrosine or L-phenylalanine synthesis. We found that glycerol served as a better carbon source than glucose, resulting in 940±46 mg/L tyramine from 4% glycerol. Next, the genes of lactate dehydrogenase (*ldhA*), formate acetyltransferase (*pflB*), phosphate acetyltransferase (*pta*), and alcohol dehydrogenase (*adhE*) were deleted to mitigate the fermentation byproduct formation. The tyramine level was further increased to 1.965±0.205 g/L in shake flasks, corresponding to 2.1 times improvement compared with that of the parental strain. By using a similar strategy, we also managed to produce 703±21 mg/L dopamine and 555±50 mg/L phenethylamine. In summary, we have demonstrated that the knockout of *ldhA*-*pflB*-*pta*-*adhE* is an effective strategy in improving aromatic amine productions, and achieved the highest aromatic amine titers in *E. coli* under shake flasks reported to date.

**Key points:** Aromatic amino acid decarboxylase from *E. faecium* was used for aromatic amine production; *ldhA, pflB, pta* together with *adhE* were deleted to mitigate the fermentation byproduct formation; Our work represented the best aromatic amine titers reported in *E. coli* under shake flasks.

## Introduction

Aromatic compounds represent a large and diverse class of chemicals that are widely used in manufacturing solvents, polymers, fine chemicals, feed and food additives, nutraceuticals, and medicines (Huccetogullari et al. 2019; Shen et al. 2020; Wang et al. 2018). Among them, aromatic amines with diverse physical characteristics are often employed as antioxidants, and precursors to pharmaceutical products (Masuo et al. 2016; Minami 2013). For example, tyramine is a high-value industrial product with widespread applications in medicine (Beltran et al. 2011). Dopamine acts as an intermediate in the biosynthesis of epinephrine and other drugs (Davie 2008; Jeong et al. 2018), and also can influence the physiological activity of plants and humans (Liang et al. 2018; Wise 2004). In addition, phenylethylamine is a precursor of antidepressants for addiction cessation (Brackins et al. 2011; Dwoskin et al. 2006). To date, the biomanufacturing process of aromatic amines mainly relies on chemical synthesis (Corrigan et al. 1945; Epstein et al. 1964). However, the chemical method typically involves complicated steps, harsh reaction conditions, and non-renewable feedstock, which is considered an environment-unfriendly process.

With the fast advancements of synthetic biology and metabolic engineering, microbial synthesis of chemicals has made significant strides in recent years (Cho et al. 2015; Huccetogullari et al. 2019). For instance, *Escherichia coli* has been extensively utilized as the host for synthesizing many natural products, owing to its well-characterized genetic information and abundant molecular tools (Yang et al. 2020). Aromatic amines typically use aromatic amino acids as precursors, which are synthesized primarily through the shikimate pathway (SHK) in microorganisms (Averesch and Kromer 2018; Shen et al. 2020). The SHK pathway is initiated by the condensation of phosphoenolpyruvate (PEP) from the glycolytic pathway and D-erythrose 4-phosphate (E4P) in the pentose phosphate (PP) pathway to produce 3-deoxy-D-arabino-heptulosonate-7-phosphate (DAHP), which is subsequently transformed to aromatic amino acids (Averesch and Kromer 2018; Cao et al. 2020) (Fig. 1). Koma *et al*. overexpressed a cluster of genes from the SHK pathway and a heterologous decarboxylase gene in *E. coli*, and the resulting strain produced 6.3 mM (863mg/L) tyramine (Koma et al. 2012). Heterologous expression of tyrosinase and decarboxylase in the L-tyrosine overproducing *E. coli* resulted in 260 mg/L dopamine (Nakagawa et al. 2011).

**Fig. 1.**
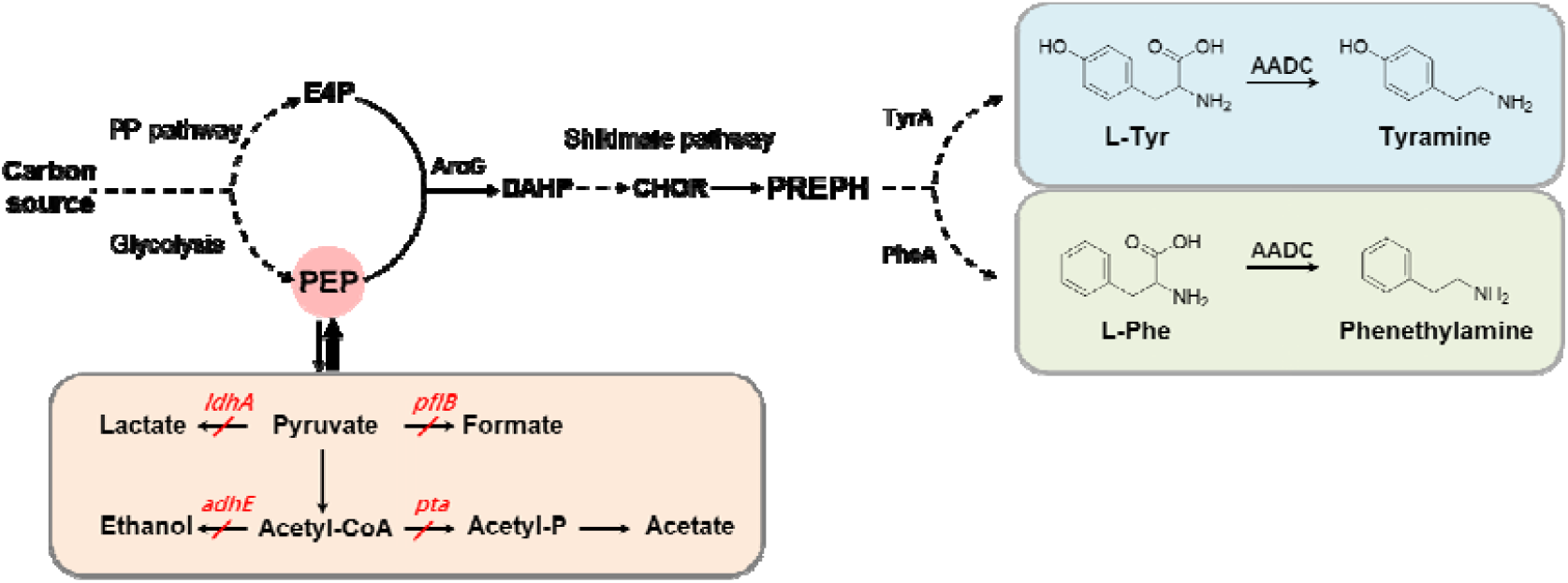
Schematic diagram of aromatic amine production from the shikimate (SHK) pathway. PP pathway: pentose phosphate pathway; AroG, 3-deoxy-D-arabinoheptulosonate-7-phosphate synthase; TyrA, chorismate mutase/prephenate dehydrogenase; PheA, chorismate mutase/prephenate dehydratase; HapBC, 4-hydroxyphenylacetate 3-monooxygenase; AADC, aromatic amino acid decarboxylase. DHAP: Dihydroxyacetone phosphate; E4P: D-erythrose 4-phosphate; PEP: Phosphoenolpyruvate; DAHP: 3-deoxy-D-arabino-heptulosonate-7-phosphate; CHOR: Chorismate; PREPH: Prephenate; L-Tyr, L-tyrosine; L-Phe, L-phenylalanine. *ldhA* encodes D-lactate dehydrogenase; *pflB* encodes pyruvate formate lyase; *pta* encodes phosphotransacetylase; *adhE* encodes alcohol dehydrogenase. The red box highlights the fermentaion byproduct related pathways. Dashed lines illustrate multiple steps.

In this work, we aimed to develop an *E. coli* platform to improve the biosynthesis of aromatic amines. As shown in Fig. 1, the main strategy used to improve aromatic amines production comprises the deletion of fermentation byproduct-related pathways to enhance the metabolic flux of the SHK pathway. In particular, we investigated the effect on aromatic amine biosynthesis by knockout of lactate dehydrogenase (*ldhA*), formate acetyltransferase (*pflB*), phosphate acetyltransferase (*pta*), and alcohol dehydrogenase (*adhE*).

## Materials and methods

### Strains and reagents

*E. coli* DH5α was utilized to construct plasmids, and *E. coli* MG1655 (DE3) derived strain with ΔtyrA and ΔpheA (Lai et al. 2022) was employed as the chassis cell for aromatic amine production. LB medium containing 10 g/L tryptone, 5 g/L yeast extract, and 10 g/L NaCl was applied for cultivating *E. coli*. Appropriate antibiotics (100 μg/mL ampicillin, 50 μg/mL kanamycin, 34 μg/mL chloramphenicol) were supplemented to maintain the plasmids when needed. Enzymes (high-fidelity phusion polymerase, *Bam*HI-HF, *Xho*I, *Bsa*I-HF, *Esp*3I-HF, and T4 DNA ligase) were purchased from New England Biolabs (Beverly, MA, USA). PCR purification kit, gel extraction kit, and plasmid DNA extraction kit were all purchased from BioFlux (Shanghai, China). The details of chemicals used in this study are provided in Supplementary materials.

### Plasmid and strain construction

All the genes were PCR amplified using high-fidelity phusion polymerase. The oligonucleotides used in this study are listed in Supplementary Table S1. The gene encoding *pheDC* from *E. faecium* (Genbank: AJ783966.1) was codon optimized for *E. coli* and synthesized by GenScript (Nanjing, China). The genes encoding *hpaBC* (Genbank: Z37980) were obtained from the genomic DNA of *E. coli*.

The plasmid pET-AroG^fbr^-TyrA^fbr^ and pET-AroG^fbr^-PheA^fbr^ were constructed in a similar way to our previous report (Lai et al. 2022). In brief, the genes encoding *tyrA*^*fbr*^, *pheA*^*fbr*^, and *aroG*^*fbr*^ were obtained from the genomic DNA of *E. coli* via overlapping PCR amplification process, digested with *Esp*3I, and ligated into pETDuet-1 between *Bam*HI and *Xho*I sites. For the plasmid pRSF-PheDC, the synthesized *pheDC* gene was inserted into pRSFDuet-1 between *Bam*HI and *Xho*I sites. For the plasmid pACYC-HpaBC, the *hpaBC* gene was amplified using *E. coli* genomic DNA as a template, and inserted into pACYCDuet-1 between *Bam*HI and *Xho*I sites. For constructing MG1655 (DE3) derived ΔpflB-ΔldhA-Δpta-ΔadhE strain, the gene knockout procedure was carried out via the CRISPR/Cas9 method (Cong et al. 2013). The engineering strains with corresponding plasmids were obtained by standard electroporation or heat-shock approach. All the details of plasmids and strains are provided in Supplementary Table S2.

### Shake flask cultivation

Colonies were inoculated from solid agar plates into 10 mL test tubes containing 2 mL LB medium and cultivated at 37°C and 250□rpm to prepare the seed culture. The next day, fresh overnight cultures (0.15 mL) were inoculated into 50 mL shake flasks containing 15 mL modified M9 medium with appropriate antibiotics. The components of the modified M9 medium were given in Supplementary Material. Upon the cell density reaching 0.4-0.6, isopropyl β-D-1-thiogalactopyranoside (IPTG) was added to the media to a final concentration of 10 μM for inducing the gene expressions. The cell cultures were shifted to 30°C and 250□rpm for the aromatic amine productions. Samples were periodically taken for monitoring the cell growth by using a microplate reader (Biotek, Synergy H1).

### HPLC analysis of aromatic amine levels

The samples were centrifuged at 14,000□rpm for 10□min to remove the cellular pellets. Shimadzu LC-20A system equipped with a photodiode array detector and a reversed-phase C18 column (150 mm × 4.6 mm × 5 μm) was used for the quantitation of aromatic amines. The column was maintained at a temperature of 40 °C. To identify tyramine and phenethylamine, the mobile phase comprising 90% ultrapure H_2_O (supplemented with 0.1% trifluoroacetic acid) and 10% acetonitrile was used. For dopamine detection, the mobile phase containing 95% ultrapure H_2_O (supplemented with 0.1% trifluoroacetic acid) and 5% acetonitrile was used. The flow rate was maintained at 1.0 mL/min. The wavelengths used for detecting tyramine, dopamine and phenethylamine were set at 222 nm, 203 nm, and 208 nm, respectively. The retention times of tyramine, dopamine and phenethylamine were 3.5 min, 3.8 min and 8.3□min, respectively. The aromatic amine levels were quantitated using an external standard curve based on authentic standards.

## Results

### *De novo* production of tyramine in *E. coli*

As shown in Fig.1, aromatic amines such as tyramine and phenethylamine can be produced by coupling the aromatic amino acid synthesis with heterologous expression of aromatic amino acid decarboxylase (AADC). In this study, we chose AADC from *E. faecium* (PheDC) as it has been functionally expressed in *E. coli*, resulting in L-phenylalanine and L-tyrosine decarboxylase activities (Marcobal et al. 2006). To increase the metabolic flux toward the SHK pathway, the feedback-resistant genes encoding 3-deoxy-D-arabinoheptulosonate-7-phosphate synthase (*aroG*) together with chorismate mutase/prephenate dehydratase (*pheA*) or chorismate mutase/prephenate dehydrogenase (*tyrA*) were overexpressed in MG1655 (DE3) with ΔtyrA ΔpheA. As shown in Fig.2a, we constructed two plasmids for expressing *aroG*^*fbr*^, *tyrA*^*fbr*^, and *pheDC* for tyramine production in *E. coli*. To identify suitable carbon sources for the synthesis of aromatic amines, we compared the modified M9 media with 4% (w/v) glycerol or glucose for tyramine productions. As shown in Fig. 2b, the tyramine titer reached 940±46 mg/L when 4% glycerol was used, whereas only 656±72 mg/L tyramine was obtained in glucose-containing medium. The maximum levels of tyrpine were achieved around 36 h, and further cultivation did not obviously improve the titer.

**Fig. 2.**
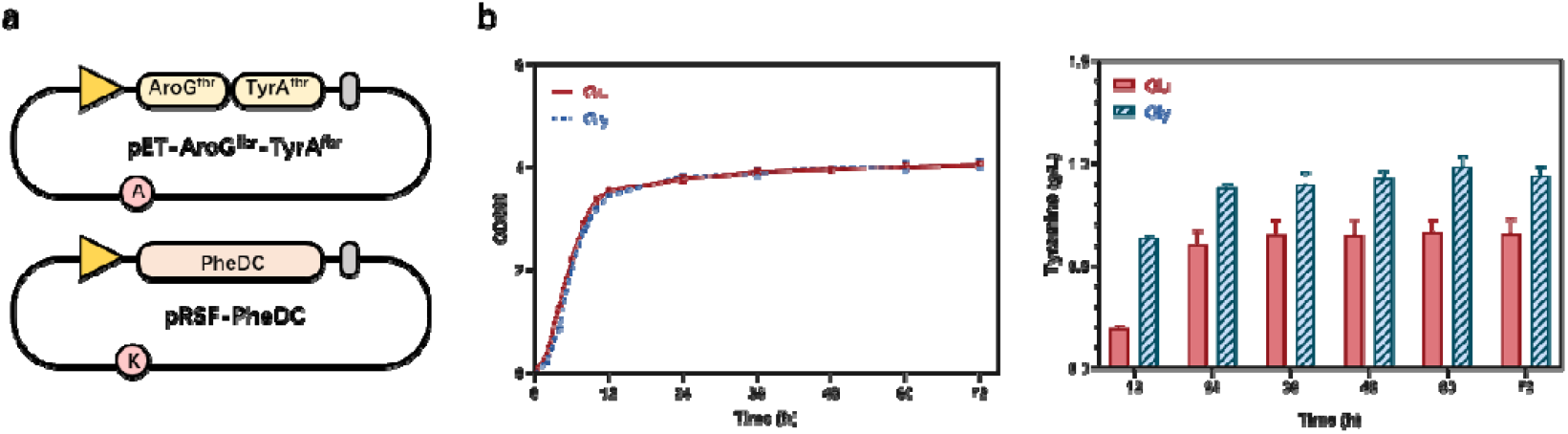
Engineering *E. coli* for tyramine overproduction. (**a**) The plasmids for tyramine production. *AroG*^*fbr*^, feedback resistant 3-deoxy-d-arabinoheptulosonate-7-phosphate synthase; *TyrA*^*fbr*^, feedback resistant chorismate mutase/prephenate dehydrogenase; *PheDC*, aromatic amino acid decarboxylase from *E. faecium*. (**b**) Growth profile and tyramine production of strain TA1.0 under shake flasks. All the experiments were carried out in 15 mL modified M9 medium containing 40 g/L glycerol or 40 g/L glucose. Experiments were performed in triplicate biological repeats and the data represent the mean value with standard deviation.

### Tyramine production by abolishing fermentation side-pathways

When cultured in the absence of oxygen, *E. coli* undergoes a mixed acid fermentation (Forster and Gescher 2014), resulting in the production of ethanol, acetate, lactate, and formate (Clark 1989; Gonzalez et al. 2008). These fermentation byproducts are respectively mediated by lactate dehydrogenase (*idhA*) and formate acetyltransferase (*pflB*) in the pyruvate catabolism and by phosphate acetyltransferase (*pta*) and alcohol dehydrogenase (*adhE*) in the acetyl-CoA catabolism (Clark 1989; Trotter et al. 2011). Even under aerobic circumstances, *E. coli* diverts a significant amount of carbon flow to fermentation byproducts as a consequence of glycolytic overflow (Kang et al. 2009). It was reported that the accumulation of by-products such as lactate, acetate, and formate would arrest the cell growth (Causey et al. 2004). In addition, James Liao’s group has demonstrated that the knockout of genes that contribute to the fermentation byproduct formation such as *adhE, ldhA, pflB*, and *pta* could substantially improve isobutanol productions in *E. coli* (Atsumi et al. 2008). Therefore, the reduced formation of fermentation byproducts would theoretically channel more carbon flux to the products-of-interest, and maximize the productivity. In this study, we further proceeded to engineer the *E. coli* metabolism by mitigating byproduct formations, so that more metabolic flux could be diverted to aromatic amines. In particular, lactate dehydrogenase (*ldhA*), formate acetyltransferase (*pflB*), phosphate acetyltransferase (*pta*), and alcohol dehydrogenase (*adhE*) were deleted by CRISPR/Cas9 mediated approach (Fig. 3a). According to Fig. 3b, the deletions of side pathway genes resulted in a slightly slower growth rate of strain TA3.0 than that of TA1.0 at the initial stage, indicating that disruption of fermentation related pathways would slightly affect the cells growth at the initial phase. However, strain TA3.0 surpassed the growth of strain TA1.0 after 12 h, probably because more carbon flux toward the TCA cycle enhances energy utilization, resulting in a higher cell density at the later phase. As shown in Fig. 3b, strain TA3.0 produced 1.965±0.205 g/L tyramine at 72 h, which is 2.1 times compared with that of strain TA1.0 (940±46 mg/L), confirming that disruption of fermentation related pathways could effectively increase more carbon flux toward aromatic amine synthesis.

**Fig. 3.**
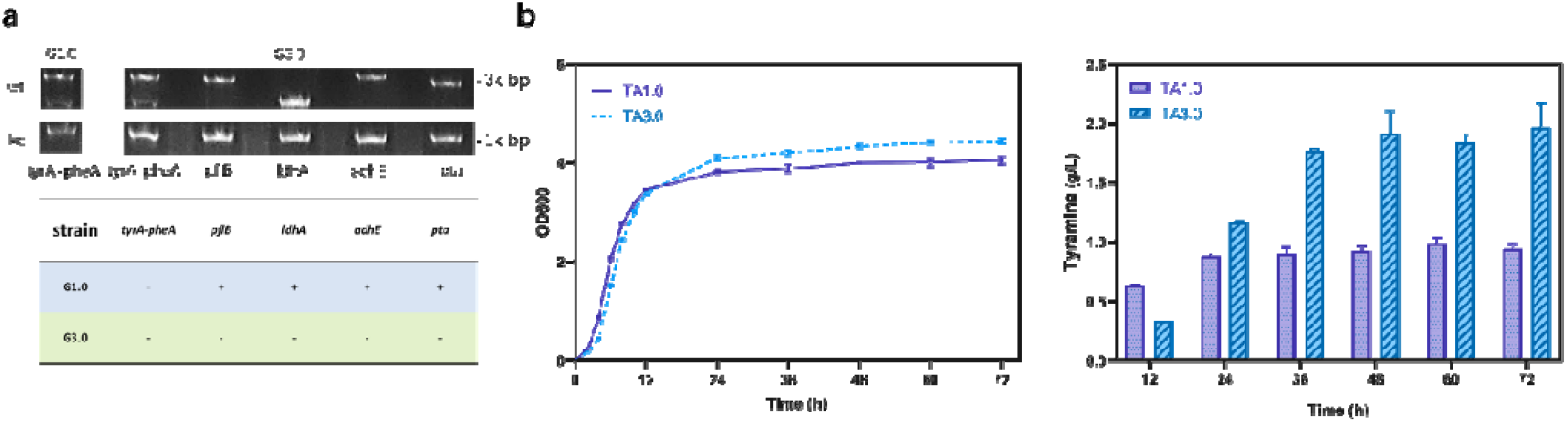
Knockout of fermentation byproduct related pathways substantially improved tyramine production. (**a**) Agarose gel image for PCR verification of gene knockout events. Strain G1.0 strain with ΔtyrA-ΔpheA was used as the control. Strain G3.0 with further deletion of *pflB, ldhA, adhE*, and *pta* was confirmed by diagnostic PCR. (**b**) Growth profile and tyramine production of strains TA1.0 and TA3.0. All the experiments were carried out in 15 mL modified M9 medium containing 40 g/L glycerol. Experiments were performed in triplicate biological repeats and the data represent the mean value with standard deviation.

### Dopamine production using the strains carrying the *hpaBC* gene

For investigating the feasibility of this platform for producing other aromatic amines, we continued our efforts to synthesize dopamine, an important pharmaceutical compound. Endogenous 4-hydroxyphenylacetate 3-monooxygenase from *E. coli* is an enzyme with two components encoded by *hpaB* and *hpaC* genes that adds a second hydroxyl group at the *ortho* position to 4-hydroxyphenylacetate and L-tyrosine (Guo et al. 2021). As shown in Fig. 4a, L-tyrosine can be hydroxylated to form L-dihydroxyphenylalanine (L-DOPA) by HpaBC, which is further decarboxylated to dopamine. Alternatively, the hydroxylation step might also occur at 4-hydroxyphenylacetate or tyramine. Overall, the enzyme cascade for dopamine production comprises AroG^fbr^, TyrA^fbr^, HpaBC, and PheDC (Fig. 4b). As shown in Fig. 4c, there was no significant difference in cell growth between strain DA1.0 and DA3.0 during dopamine synthesis. Notably, strain DA3.0 exhibited the highest level of dopamine production (703±21 mg/L) at 48 h, which is nearly 2.9 times than that of stain DA1.0 (242±24 mg/L). Therefore, we achieved a considerable improvement in dopamine production than the previous study (Nakagawa et al. 2011).

**Fig. 4.**
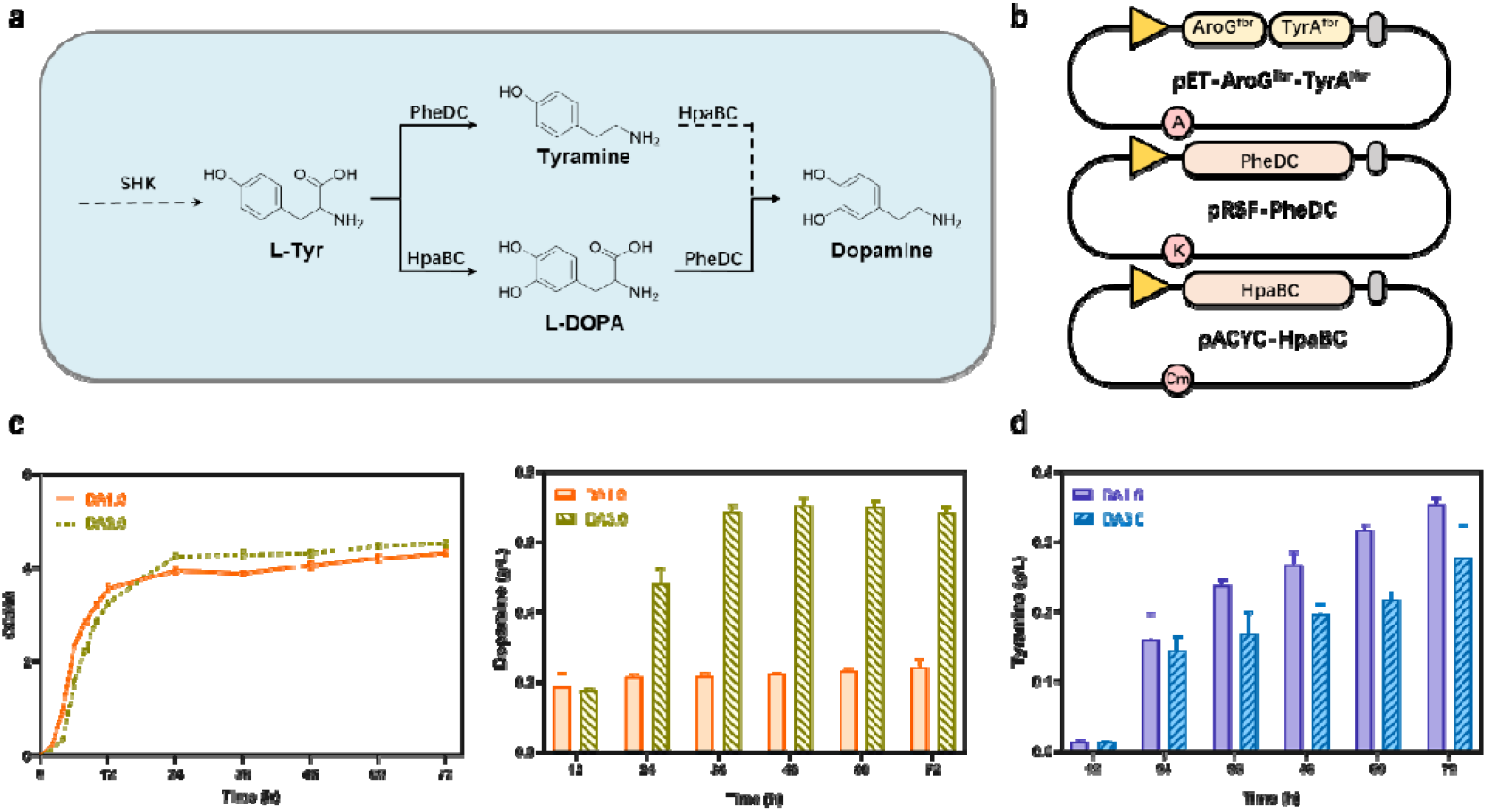
The dopamine production in shake flasks. (**a**) The proposed biosynthetic route toward dopamine synthesis. The dashed arrow of HpaBC indicates the poor activity of HpaBC in converting tyramine to dopamine. (**b**) The plasmids for dopamine production. (**c**) Growth profile and dopamine production of strains DA1.0 and DA3.0. (**d**) Tyramine accumulation in strains DA1.0 and DA3.0. All the experiments were carried out in 15 mL modified M9 medium containing 40 g/L glycerol. Experiments were performed in triplicate biological repeats and the data represent the mean value with standard deviation.

As mentioned above, L-tyrosine can also be first converted to tyramine under the action of PheDC (Fig. 4a). To this end, we also measured the accumulation of tyramine during dopamine production. As shown in Fig. 4d, both strain DA1.0 and DA3.0 accumulated a substantial amount of tyramine, indicating the HpaBC activity toward tyramine hydroxylation might be not sufficient. Surprisingly, more tyramine was accumulated in the DA1.0 strain than that of strain DA3.0. We reasoned that the hydroxylation reaction requires a large amount of NADH and NADPH, critical cofactors for the flavin reduction by *hpaC*. Therefore, knockout of side-pathways that consume NADH and NADPH might favor the hydroxylation reaction with improved dopamine production in strain DA3.0.

### Phenylethylamine production using the engineered *E. coli* strain

Next, we also proceeded with phenylethylamine production in the engineered *E. coli*. Briefly, we replaced *aroG*^*fbr*^-*tyrA*^*fbr*^ plasmid with *aroG*^*fbr*^-*pheA*^*fbr*^ to switch the metabolic flux from L-tyrosine to L-phenylalanine synthesis. The two-plasmid system with enzyme cascade including *aroG*^*fbr*^*-pheA*^*fbr*^ and *pheDC* was used to convert L-phenylalanine to phenylethylamine (Fig. 5a). As depicted in Fig. 5b, the maximum phenethylamine production of 555±50 mg/L was achieved by strain PEA3.0 after 72 h, which was increased by 2.28-fold compared with that of strain PEA1.0 (169±20 mg/L). The phenethylamine level achieved in the shake flask by our study was also much higher than that of a recent report (Xu and Zhang 2020). In addition, we found that both strain PEA1.0 and PEA3.0 gave a similar growth profile (Fig. 5b). However, the final cell densities for phenylethylamine-producing strains were slightly lower than dopamine or tyramine-producing strains. Since strain PEA1.0 and 3.0 with different levels of phenylethylamine gave a similar growth rate, we concluded that phenylethylamine is not toxic to the cells at the current concentrations. Therefore, it is likely that the reduced biomass of phenylethylamine-producing strains was mainly caused by Δ*tyrA*.

**Fig. 5.**
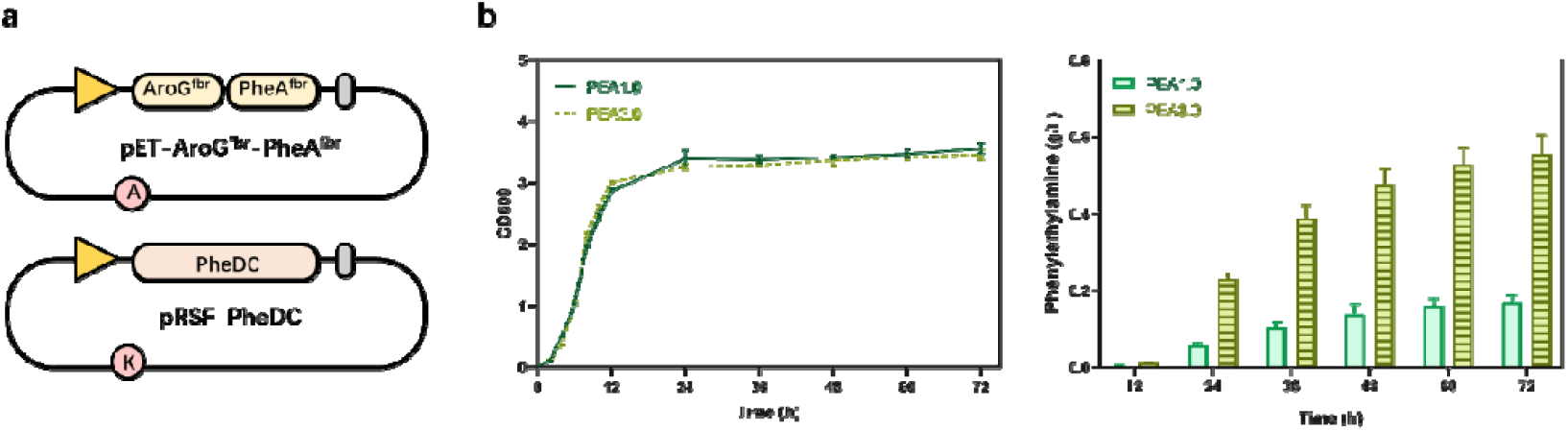
The phenethylamine production in shake flasks. (**a**) The plasmids for phenethylamine production. PheA^fbr^, feedback-resistant chorismate mutase/prephenate dehydratase. (**b**) Growth profile and phenethylamine production of strains PEA1.0 and PEA3.0. All the experiments were carried out in 15 mL of modified M9 medium containing 40 g/L glycerol. Experiments were performed in triplicate biological repeats and the data represent the mean value with standard deviation.

## Discussion

Aromatic amines have a wide range of applications in medicine, chemistry, and biology. Rapid advances in synthetic biology have facilitated the study of aromatic compound biosynthesis and offered an engineering framework for producing high-value aromatic compounds. *E. coli* has been genetically engineered to boost the productivity of aromatic compounds by removing or inhibiting undesirable genes and increasing the expression of rate-limiting genes. In this study, we applied similar strategies to increase the metabolic flux toward the SHK pathway. By introducing feedback-resistant versions of *tyrA*^*fbr*^*/pheA*^*fbr*^ and *aroG*^*fbr*^, the recombinant strains with a further expression of *pheDC* from *E. faecium* were able to produce 940±46 mg/L tyramine, 242±24 mg/L dopamine, and 169±20 mg/L phenylethylamine, respectively. According to recent studies, decreasing mixed acid fermentation is proved to be effective in increasing the production of 3-hydroxypropionate (3HP) (Liu et al. 2018), polyhydroxyalkanoate (Jung et al. 2019), β-alanine (Zou et al. 2020), and 2,3-butanediol (2,3-BD) (Song et al. 2019). Therefore, we further constructed a high titer aromatic amine production platform by abolishing fermentation side-pathways. Namely, the genes of (i) *ldhA* and *pflB* (from pyruvate to lactate and formate) and (ii) *pta* and *adhE* (from acetyl-CoA to acetate and ethanol) were deleted to improve the metabolic flux from the carbon source to the SHK pathway. Finally, 1.965±0.205 g/L tyramine, 703±21 mg/L dopamine, and 555±50 mg/L phenethylamine were obtained from 40 g/L glycerol in shake flask cultivation. The titers achieved by us were much higher than previous reports under shake flask conditions (Koma et al. 2012; Nakagawa et al. 2011; Xu and Zhang 2020).

During the dopamine biosynthesis, we observed that both strain DA1.0 and DA3.0 accumulated a substantial amount of tyramine, suggesting that the HpaBC could not effectively hydroxylate tyramine to dopamine. Considering that a mutant HpaBC was recently identified with good activity toward the tyramine hydroxylation (Chen et al. 2019), it will be possible to address tyramine accumulation by simply introducing the mutant HpaBC to our engineered *E. coli* strain. In addition, we were surprised to find out that less tyramine was accumulated in strain DA3.0 when compared to that of strain DA1.0. Since the hydroxylation reaction requires additional reducing power for flavin recycling, the knockout of fermentation side-pathways that consume NADH and NADPH would also improve dopamine synthesis. Therefore, it is likely that our engineered platform would be of great interest for hosting other biosynthetic pathways that require cofactors such as NADH and NADPH.

In summary, we have optimized the SHK pathway and eliminated the side-pathways involved in the mixed acid fermentation to enable high-titer generation of aromatic amines in metabolically modified *E. coli*. We demonstrated that the knockout of *ldhA*-*pflB*-*pta*-*adhE* is an effective strategy in improving aromatic amine productions. Based on our findings, we believe that the *E. coli* system has great prospects for the future industrial-scale aromatic amine production. Moreover, these engineered strains might also be used to manufacture other aromatic compounds with pharmaceutical value.

## Author contributions

J. Y. conceived and designed the project. T. Y. and P. W. constructed the plasmids, strains and collected the data. T. Y. and Y. Z. analyzed the date. T.Y., Y. Z. and J. Y. wrote the manuscript.

## Compliance with ethical standards

### Ethical approval

This study does not contain any studies with human participants or animals performed by any of the authors.

### Data availability

All data generated or analyzed during this study are included in this published article [and its supplementary information files].

---

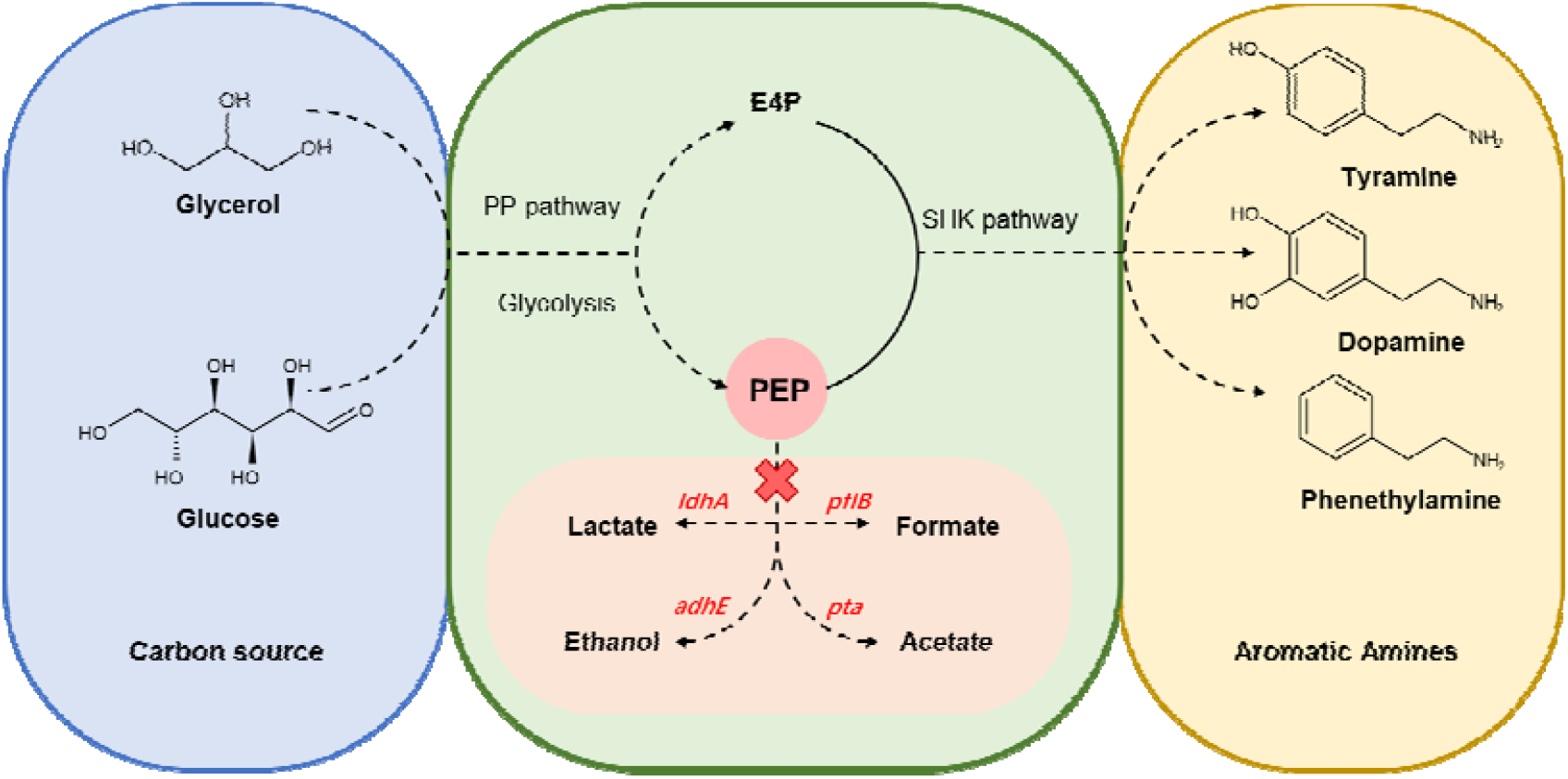
TOC

